# *Wolbachia, Cardinium* and climate: an analysis of global data

**DOI:** 10.1101/490284

**Authors:** J. Charlesworth, L. A. Weinert, E. V. Araujo-Jnr., J. J. Welch

## Abstract

Bacterial secondary symbionts are very common in terrestrial arthropods, but infection levels vary widely among populations. Experiments and within-species comparisons both suggest that environmental temperature might be important in explaining this variation. To investigate the importance of temperature, at broad geographical and taxonomic scales, we extended a global database of terrestrial arthropods screened for *Wolbachia* and *Cardinium*. Our final data set contained data from 114,297 arthropods (>2,500 species) screened for *Wolbachia* and 17,011 arthropods (>800 species) screened for *Cardinium*, including population samples from 137 different countries, and with mean temperatures varying from - 6.5 to 29.2°C. In insects and relatives, *Cardinium* infection showed a clear and consistent tendency to increase with temperature. For *Wolbachia*, a tendency to increase with temperature in temperate climates, is counteracted by reduced prevalence in the tropics, resulting in a weak negative trend overall. We discuss the implications of these results for natural and introduced symbionts, in regions affected by climate change.

## Introduction

Bacterial endosymbionts are common in all major groups of arthropods, and can exert profound effects on their hosts (Zchori-Fein and Bourtzis 2011, Duron & Hurst 2013, Weinert et al. 2015). While infection is very widespread, symbiont prevalence (i.e., the proportion of individuals infected) varies widely among populations. Several factors might shape this variation, including the costs of reproductive parasitism (Hurst and Frost 2015), the benefits of protection against viral pathogens (Moreira et al 2008), or host dispersal patterns (Jiggins 2017).

Another putative influence on endosymbiont prevalence is environmental temperature (Corbin et al 2017). Laboratory studies have suggested that endosymbionts are more susceptible to thermal stress than their hosts (Wernegreen 2012; Kikuchi et al. 2016; Hussain et al. 2017), and the physiological costs or benefits of carriage, or endosymbiont transmission might also be temperature-sensitive (Corbin et al 2017). Evidence of a role for temperature also comes from comparative biogeography. For example, *Cardinium* infection in *Culicoides* midges is higher in warmer areas of Israel (Morag et al. 2012), while similar trends have been reported for *Wolbachia* in *Lepidoptera* and *Diptera* (Ahmed et al. 2015; Morrow et al. 2015; Kreisner et al. 2016). However, these remain isolated results, and sample prevalence in *Wolbachia* is remarkably constant at the broadest continental scales (Werren and Windsor 2000).

Here, we survey global data on symbiont prevalence in wild populations of terrestrial arthropods. We focus on *Wolbachia* and *Cardinium*, which are well sampled, and infect, respectively, around 1/2 and 1/8 of all terrestrial species (Weinert et al. 2015). We then test for a relationship between prevalence and the climate of the sampling location.

## Results

We collated data from 320 publications, comprising 135,875 individual arthropods screened. When sampling location was specified, we then obtained an estimate of mean temperature between 1970-2000 (Fick and Hijmens 2017), and a categorical assignment of climate, using the Köppen system, which summarizes multiple ecologically relevant variables, such as seasonal precipitation and heat (Peel et al 2007; Chen and Chen 2013). The data are diverse taxonomically (with hosts from >45 arthropod orders), and geographically (with 27/31 Köppen climates, including all five higher-level classifications: tropical, arid, temperate, continental, and polar). The full database is available as Supplementary Table S1, and is summarized in Figures 1-2, and S1-S3.

**Figure 1:**
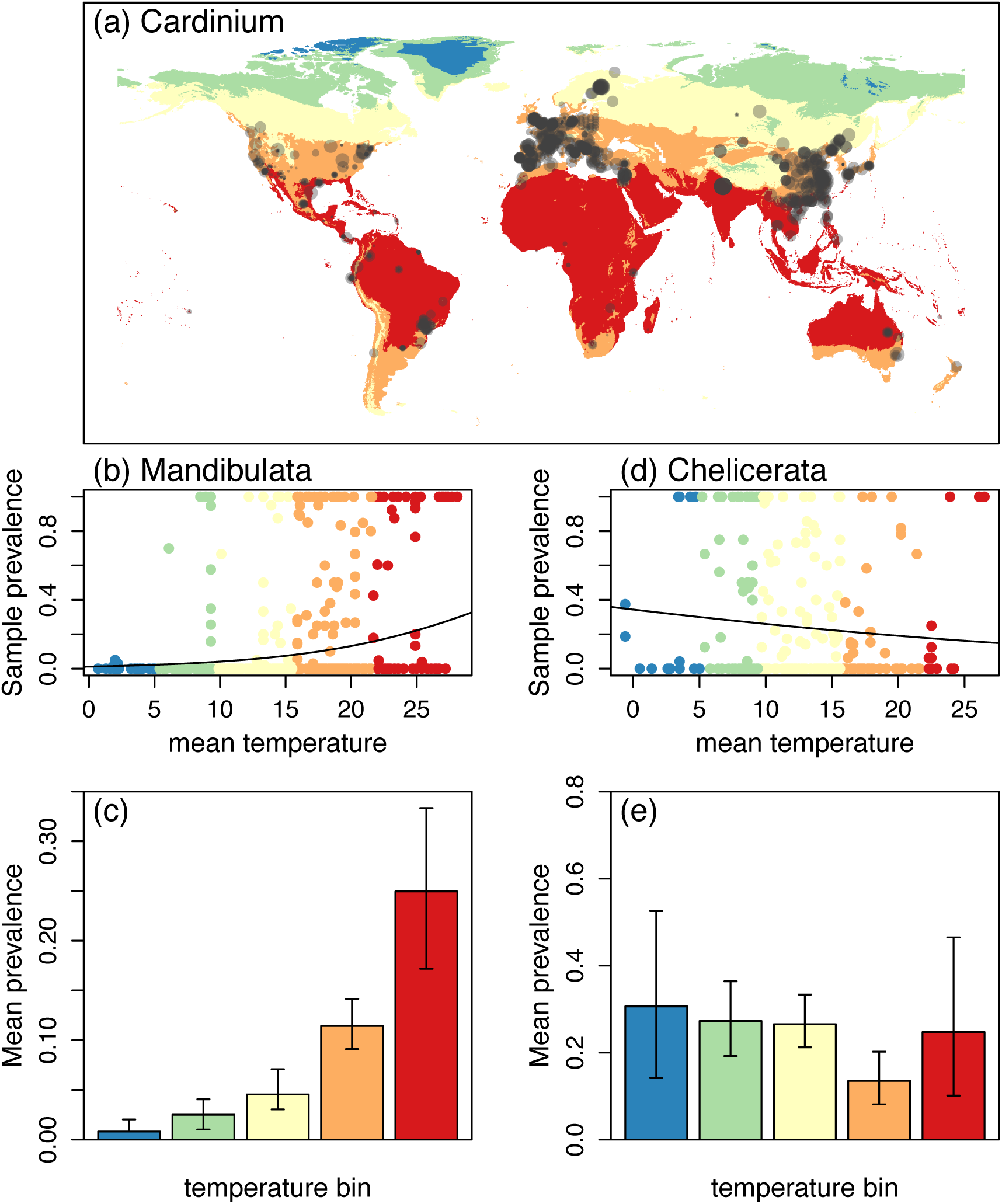
Climatic variation in the prevalence of *Cardinium* infection in terrestrial arthropods. (a): world map, with color corresponding to mean temperature over the period 1970-2000. Points indicate population screens included in the present study, with point size indicating the number of individuals sampled; (b) and (d): regression of mean prevalence (logit transformed) on the mean temperature of sampling location; the best-fit lines correspond to the three-parameter model reported in Table 1. (c) and (e): illustrative plots, showing the estimated mean prevalence for populations falling within a temperature range, as indicated by color in the panels above. Separate results are given for the two major clades of arthropods, which have very different levels of *Cardinium* infection, namely (b)-(c) Mandibulata (mostly insects, but also wingless hexapods, crustaceans and myriapods), and (d)-(e) Chelicerata (ticks, mites, spiders and relatives).

We first consider results for *Cardinium* (Figure 1). This symbiont has very different rates of infection in the two major arthropod groups, namely Chelicerata: mites, ticks, spiders and relatives, and Mandibulata: mainly insects, but also wingless hexapods, crustaceans and myriapods (Weinert et al. 2015; Martin and Goodacre 2012), and so we consider these two major groups separately.

**Table 1:** The effects of mean temperature and climatic zone on symbiont prevalence

| **Data set:** | KC | *np* | AIC | temp: slope (*p* val) |  |
| --- | --- | --- | --- | --- | --- |
| **(a) *Cardinium* Mandibulata** | - | 2 | 1446.19 |  |  |
| # Populations: 1376 | - | 3 | 1396.30 | 0.127 | (9.19×10<sup>-13</sup>*) |
| # Individuals (infected): 11,757 (2072) | K5 | 6 | 1445.05 |  |  |
| # Species min-max: 627-807 | K5 | 7 | 1370.94 | 0.214 | (4.09×10<sup>-17</sup>*) |
|  | K31 | 20 | 1380.66 |  |  |
|  | K31 | 21 | **1349.59** | 0.192 | (6.74×10<sup>-9</sup>*) |
| **(b) *Wolbachia* Mandibulata** | - | 2 | 22039.02 |  |  |
| # Populations: 7,988 | - | 3 | 22028.35 | -0.011 | (0.000368*) |
| # Individuals (infected): 102,269 (47,734) | K5 | 6 | 21976.81 |  |  |
| # Species min-max: 2,461-3,693 | K5 | 7 | 21958.93 | 0.024 | (7.78×10<sup>-6</sup>*) |
|  | K31 | 27 | **21747.23** |  |  |
|  | K31 | 28 | 21748.92 | 0.004 | (0.572) |
| **(c) *Cardinium* Chelicerata** | - | 2 | 973.90 |  |  |
| # Populations: 343 | - | 3 | **972.18** | -0.040 | (0.054) |
| # Individuals (infected): 4,716 (1,210) | K5 | 6 | 975.68 |  |  |
| # Species min-max: 137-155 | K5 | 7 | 974.42 | -0.045 | (0.070) |
|  | K31 | 21 | 974.33 |  |  |
|  | K31 | 22 | 976.20 | -0.012 | (0.711) |
| **(d) *Wolbachia* Chelicerata** | - | 2 | 2000.97 |  |  |
| # Populations: 636 | - | 3 | 2002.84 | -0.005 | (0.719) |
| # Individuals (infected): 8,410 (2,146) | K5 | 6 | 1992.71 |  |  |
| # Species min-max: 322-348 | K5 | 7 | 1994.69 | 0.003 | (0.868) |
|  | K31 | 20 | **1944.81** |  |  |
|  | K31 | 21 | 1945.09 | 0.033 | (0.181) |

*Note:* KC: climatic zones included in the model as categorical predictors, either K5 (five higher-level Köppen classifications) or K31 (up to 31 finer-grained climates); *np*: the total number of parameters in the model fit; AIC: Akaike Information Criterion (Akaike 1974), with the preferred model shown in bold; temp: the best-fit slope and *p*-value associated with the mean temperature, when this was included in the model. \* *p* < 0.005. The number of arthropod species in each subset of the data is given as a minimum (counting named species only), and a maximum (including each partially identified taxon as a unique species).

For *Cardinium* infection in Mandibulata (Figure 1a-c; Table 1a), there is a clear trend for increasing infection with temperature (Figure 1c). This is confirmed by a regression analysis (Figure 1b; Table 1a). Temperature remained a significant predictor when we also allowed prevalences to vary systematically with climatic zone, by allowing each of the five higher-level Köppen climates to have a typical prevalence level (“K5” in Table 1a). The same was also true when we allowed for systematic differences between each of the finer-grained Köppen climates (“K31” in Table 1a).

There are two possible caveats to this result. First, the best fit was obtained from the largest model (Table 1a), which might indicate model inadequacy; however, the preferred model gave a significantly better fit than equally large models, where the Köppen climate labels were permuted randomly among the sampled populations (permutation *p*-value < 10^−4^). Second, our sample of populations must be highly unrepresentative with respect to taxonomy (see Supplementary Figure S1 and Supplementary Table S2). For example, over half of the individuals sampled (6127/11,757), and three quarters of those infected (1596/2072) are Hemiptera (true bugs). Nevertheless, as we show in Supplementary Table S2, the effect of temperature remains when we remove all Hemiptera from the data set, and when we considered Hemiptera alone. Furthermore, the same trend (albeit non-significant) is evident in the two best-sampled hemipteran groups, namely sternorrhynca (aphids, whiteflies and relatives), and fulgoromorpha (planthoppers). Together, then, results suggest that both mean temperature, and some other features of climate have predictable effects on the levels of *Cardinium* infection in the Mandibulata.

For *Wolbachia*, in the same host group (Figure 2a-c; Table 1b), results are quite different. For these data, there is a tendency for colder climates to harbor higher prevalence infections (Table 1b), but the effect, though significant, is small. For example, an increase in temperature from 0° to 10° only decreases the expected mean prevalence from 42% to 40%. However, the effect does appear in multiple taxonomic groups. As shown in Supplementary Table S2, and Supplementary Figure S4, a negative effect of temperature is found in 4/6 of the well-sampled insect orders: Coleoptera, Hymenoptera, Hemiptera and Orthoptera, and in pooled data from the remaining, sparsely-sampled groups. The same trend was also seen in mosquitoes (Diptera: Culididae), and in the remainder of the Diptera, though not in the well-sampled Lepidoptera.

**Figure 2:**
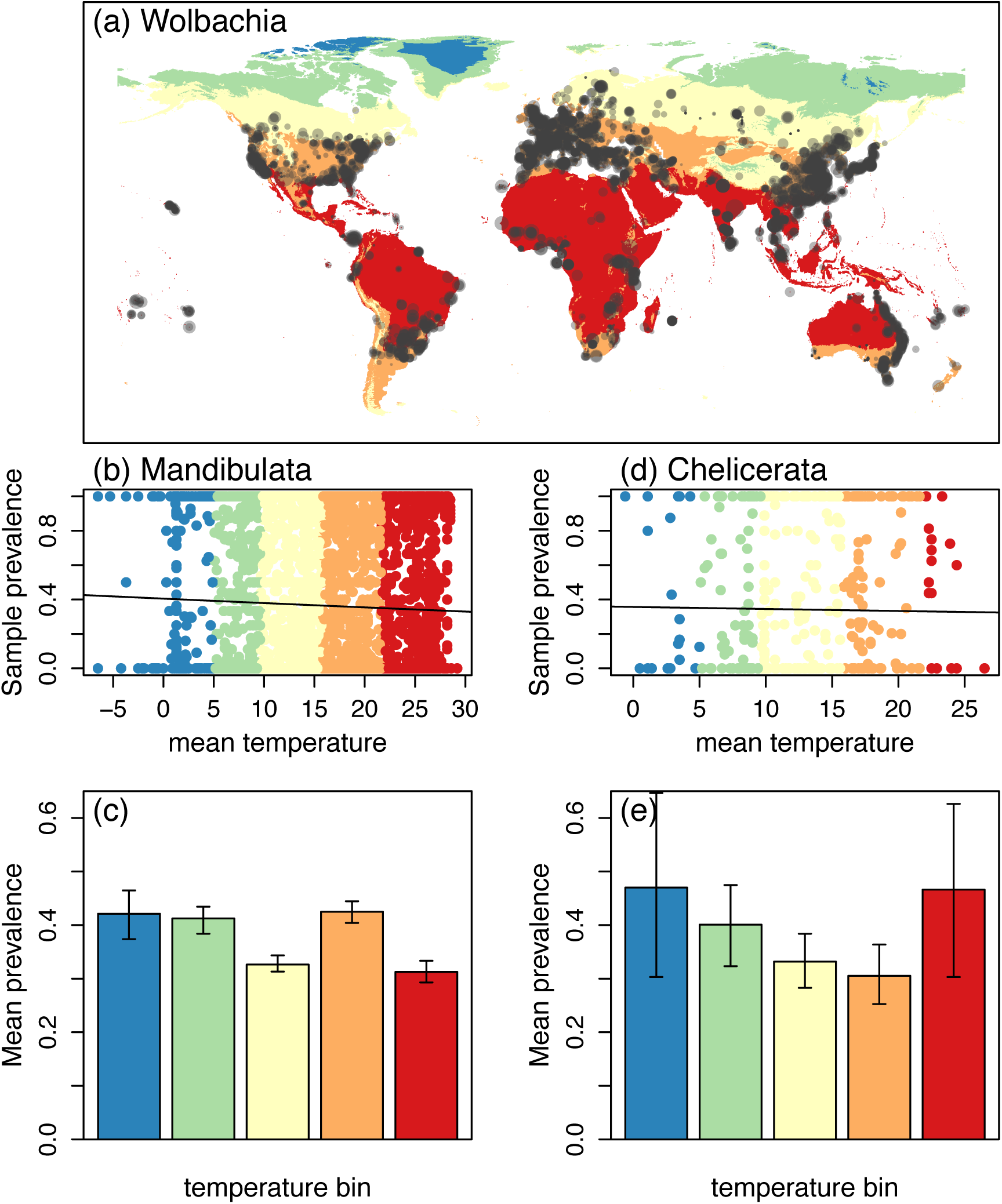
Climatic variation in the prevalence of *Wolbachia* infection in terrestrial arthropods. All details match Figure 1.

As shown in Figure 3, this effect is driven by differences between the major climatic zones, with several groups showing less infection in tropical regions, and more infection in cold, continental regions (see also Figure S9). If we model this effect, assigning a typical prevalence level to each of the five climates (“K5” in Table 1b), then the effect of temperature reverses sign, and it becomes a significantly positive predictor of prevalence. Finer-grained analyses show that this is driven by a strong effect of temperature within the best-sampled “temperate” zones, with no consistent pattern elsewhere (Supplementary Table S3 and Figure S5). The effect also disappears if we allow for systematic differences between the finer-grained Köppen climates (“K31” in Table 1b); in this case, model fit improves substantially, and more so than when climatic labels are randomly permuted (permutation *p* < 10^−4^), but including temperature as an explanatory variable adds little predictive power.

**Figure 3:**
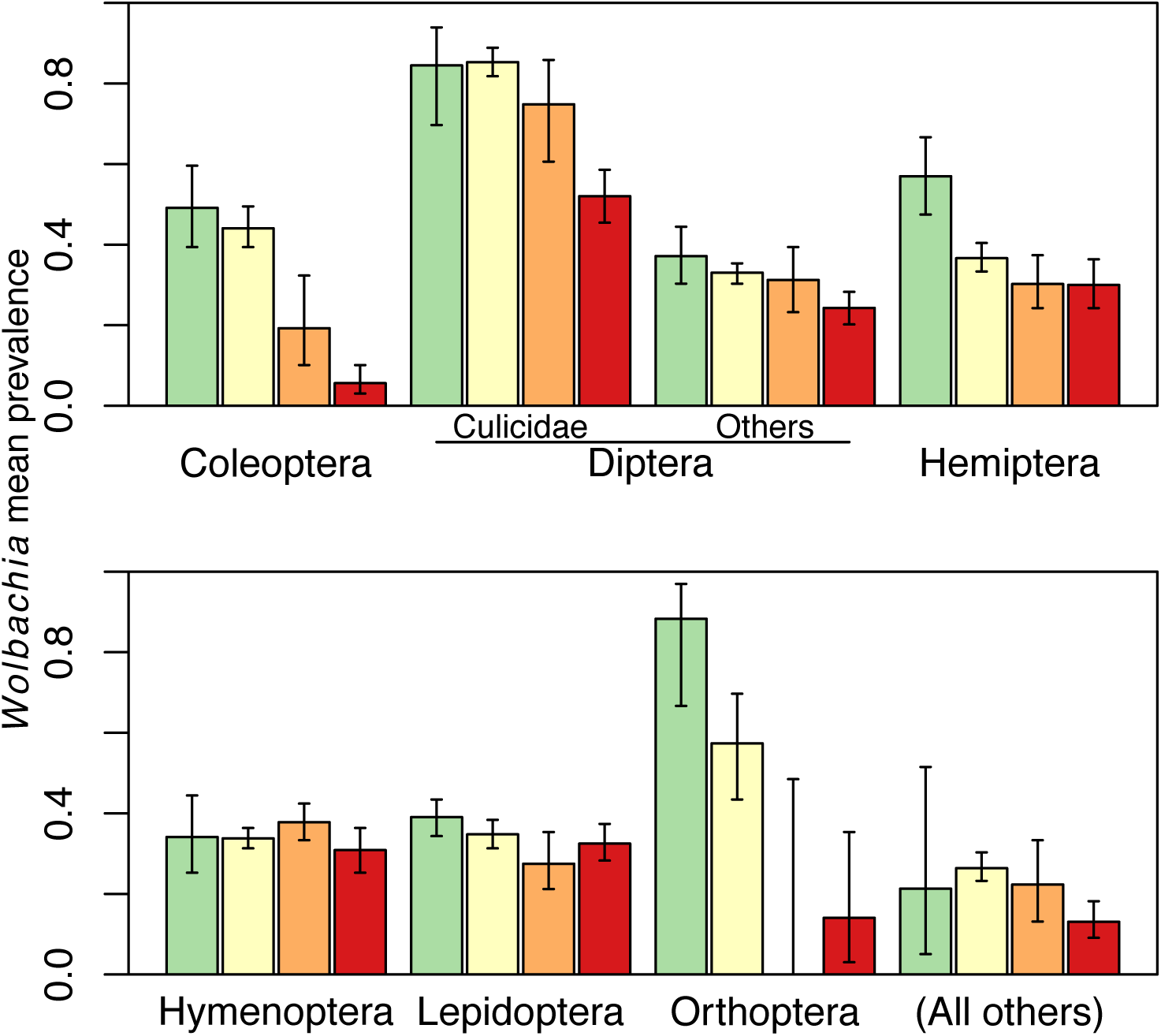
Estimated mean prevalence for *Wolbachia* infection in mosquitoes (Culicidae: Diptera), the best-sampled insect orders, and the remainder of the mandibulate arthropods. In each case, separate estimates are shown for populations from the four best-sampled climatic zones, according to the Köppen system. From left-to-right, these are Continental (green bars); Temperate (yellow bars), Arid (orange bars) and Tropical (red bars). The main database also contains samples from polar climates, but only a few for each taxonomic group.

Chelicerates are much less well sampled than insects, but this host group showed little evidence of a consistent effect of temperature (Figure 1d-e; Figure 2d-e; Table 1c-d). Indeed, for *Wolbachia* there is a notable lack of variation in the mean prevalence estimates. We find meaningful variation between typical prevalences in the finer-grained climatic zones (permutation p<10^−4^), but the best-fit slope for temperature is close to zero (Table 1d), and confidence intervals overlap for most pairs of higher-level climates (see Supplementary Figure S6). This lack of a clear trend holds for the well-sampled Acari (mites, ticks and relatives), which includes many disease vectors, and for the remainder of the chelicerate data (Supplementary Table S2).

## Discussion

We have compiled a global database of arthropod populations screened for the bacterial endosymbionts *Cardinium* and *Wolbachia*, and used these data to test for an effect of climate, and especially temperature, on infection prevalence levels. Our data were extensive, but haphazardly sampled, and so we have focussed on patterns that are found consistently in different subsets of the data.

A clear and consistent effect of temperature was found for *Cardinium* infection in Mandibulata hosts, and particularly in insects. This suggests that the pattern reported by Morag et al. (2012; for *Culicoides* midges, collected across Israel), obtains much more generally. However, this effect was limited to *Cardinium* in insects, where incidence levels are much lower than in the other host-symbiont combinations studied here (Weinert et al. 2015).

For infection with *Wolbachia* – where incidence levels are much higher – the patterns observed are more complex. In Chelicerate hosts (including mites, ticks and relatives), no consistent trends were found, and infection levels are surprisingly constant across very different geographical regions, and host groups. In insects, by contrast, we found a tendency for infection to increase with temperature, but only within temperate climatic zones. No such trend was observed within tropical climates, but these tended to have lower infection levels overall. This last pattern runs contrary to previous studies within single species or families of Diptera (Morrow et al. 2015; Kriesner et al. 2016), but was observed here in multiple host groups, including mosquitoes, the major dipteran disease vectors.

These results for *Wolbachia* have a special interest, because this symbiont is increasingly used as a biocontrol agent, particularly for mosquito-borne human pathogens. For example, in *Aedes aegypti, Wolbachia* infection inhibits the replication of dengue, chikungunya and zika viruses, as well as malarial parasites (Moreira et al 2009; Jeffries and Walker 2016; Jiggins 2017). The use of *Wolbachia* in biocontrol has, of course, focused on tropical regions (in Australia, Brazil, Indonesia and Vietnam), where the problems of mosquito-borne disease are most acute. But the trends within temperate zones suggest that changes in temperature can have consistent effects on symbioses in regions where, because of climate change, such diseases are re-emerging (Jeffries and Walker 2016; Amos 2016). Experimental studies have shown that increases in temperature can perturb the mosquito-*Wolbachia* symbioses in the laboratory (Yixin et al. 2016; Ross et al. 2017), and our results suggest these lab results could have a strong ecological relevance.

## Methods

### Database collation

Our starting point was the collation by Weinert et al. (2015), of PCR screens of terrestrial arthropods, from 364 publications prior to 2014. From this database, we retained only screens for *Wolbachia* and *Cardinium*, from wild populations. Unlike Weinert et al. (2015), we retained screens from known outbreak areas, but removed studies where arthropod individuals were pooled before screening. We re-checked entries for all studies, and reentered data in accordance with the goals of the present study. This led to several changes in previous entries (∼0.09% of the total). Using the criteria described in Weinert et al. (2015), we then searched the literature for more recent studies, stopping at March 2018. This yielded data from an additional 125 published papers, increasing the number of *Wolbachia* screens by about a third. To associate each record with climate data, we first obtained the decimal latitude and longitude of each sampling location, either from the source publications, or by contacting the authors (see acknowledgements), or by finding named locations on Google Maps (2017). Where location information was very coarse-grained (e.g. “Texas” or “Scotland”) we used two strategies to calculate approximate coordinates. If other studies had sampled from the same location, we used the median coordinates of the sampled data for our unspecified location. If no other studies sampled from a given location, we used coordinates for the centroid of that location. These approximate methods were used for ∼15.6% of records, as noted in the database.

We then used the latitude and longitude information to obtain climatic data. First, we obtained mean temperature estimates over a 2.5-minute grid (∼4.5km at the equator) from the WorldClim database (Fick and Hijmens 2017). We associated each of our sampling locations with its closest grid entry, by Euclidean distance. We also excluded samples from some island or coastal regions, where WorldClim lacks data. Each of our temperature estimates is the mean of weekly maximum and minimum temperatures, averaged over the years 1970-2000. Where sampling dates of the arthropod populations were reported, these were also recorded in our database. But despite the volume of data, there were insufficient records to test directly for temporal change. Any discrepancy between sampling dates and temperature estimates should add noise, but not bias to our results. Estimates of temperature variability, and of the maximum and minimum temperatures during 1970-2000, correlate very strongly with the mean temperatures (Supplementary Figure S7), increasing our confidence that our chosen predictor is ecologically meaningful. For the same reason, we used only the mean temperature in our analyses, and did not add additional, highly correlated predictors. We also classified each sampling location according to the Köppen system, again using Euclidean distance, to associate each sampled point with a grid reference (Peel et al 2007, data downloaded from Chen and Chen 2013). These classifications correlate with the mean temperature estimates, and this increases the stringency of our tests, but as shown in Supplementary Figure S3, most climate types include a wide range of temperatures, and there is overlap between the temperature ranges of different climatic zones.

### Statistical analyses

Following Hilgenboecker et al. 2008 and Weinert et al. 2015, our analyses use Beta-binomial modeling. Here, the number of infected individuals in a sample is drawn from a binomial distribution, parameterized with the sample size, and true population prevalence. This true prevalence is then drawn from a Beta distribution, which is parameterized with a mean prevalence, and a correlation parameter, which describes how much of the variation in infection is distributed within versus between populations. Both parameters of the Beta distribution vary between zero and one, and so the linear model uses a logit link function. While both parameters could vary with the predictors (i.e., the temperature or ecology), we are most interested in mean prevalence, and in the main text, we assume that the correlation parameter takes a constant value, also estimated from the data. As shown in Supplementary Table S2 and Supplementary Figures S8-S11, relaxing this assumption had little qualitative effect on the results. This agrees with simulation results, which suggest that Beta-binomial models yield robust estimates of the mean prevalence, even if the shape of the distribution is misspecified (Weinert et al. 2015). All models were fit using the *vglm* function of the R package VGAM v1.0-3 (Yee 2015), or by custom scripts presented by Weinert et al. (2015). For the randomization tests, we used 10,000 random permutations of the Köppen classifications associated with each population, and the *p*-value is the proportion of these permutations where the maximized log-likelihood is at least as high as with the true classifications. Separate permutations were made for population samples comprising only a single individual, and population samples containing two or more sampled individuals. This is because single-individual screens are more likely to approximate an unbiased sample of prevalence levels, but do not provide enough information to fit both parameters of the Beta distribution (Hilgenboecker et al. 2008; Weinert et al. 2015). Following Benjamin et al. (2017), we define statistical significance as *p*<0.005. For the illustrative plots, shown in Figures 1-2 panels (c) and (e), and Supplementary Figure S4, we binned populations by mean temperature, centering the bins on the mean temperatures recorded in our data set for the five higher-level Köppen climates (Supplementary Figure S3). A distinct Beta-binomial model was then fit to all populations within each bin, with confidence intervals defined as mean prevalence values that reduce the maximized log likelihood by two units (Edwards 1992). The same approach was taken for estimating the typical prevalences in each higher-level climatic zone (see Figures 3 and Supplementary S6).

## Supporting information

## References

Ahmed MZ, Araujo-Jnr EV, Welch JJ, Kawahara AY. 2015. Wolbachia in butterflies and moths: geographic structure in infection frequency. Front Zool. 12:16.

Akaike, H 1974. A new look at the statistical model identification. IEEE Transactions on Automatic Control, 19: 716–723, doi:10.1109/TAC.1974.1100705

Amon JJ 2016. The impact of climate change and population mobility on neglected tropical disease elimination. International Journal of Infectious Diseases 53:12.

Benjamin, D. J., J. Berger, M. Johannesson, B. A Nosek, E.-J. Wagenmakers, R. Berk, K. Bollen, et al. 2017. “Redefine Statistical Significance”. PsyArXiv. July 22. psyarxiv.com/mky9j.

Chen, D. and H. W. Chen, 2013: Using the Köppen classification to quantify climate variation and change: An example for 1901–2010. Environmental Development, 6, 69-79, 10.1016/j.envdev.2013.03.007.

Corbin C, Heyworth ER, Ferrari J, Hurst GD (2017). Heritable symbionts in a world of varying temperature. Heredity 118:10–20.

Duron O, Hurst GD 2013. Arthropods and inherited bacteria: from counting the symbionts to understanding how symbionts count. BMC Biology 11:45. doi:10.1186/1741-7007-11-45.

Edwards, AWF. 1992. Likelihood: Expanded edition. The Johns Hopkins University Press, Oxford.

Fick SE and Hijmans RJ. 2017. WorldClim 2: new 1-km spatial resolution climate surfaces for global land areas. Int. J. Climatol, 37: 4302–4315. doi:10.1002/joc.5086

Hilgenboecker K, Hammerstein P, Schlattmann P, Telschow A, Werren JH 2008. How many species are infected with Wolbachia? - a statistical analysis of current data. FEMS Microbiol Lett. 281(2):215–20.

Hurst GD, Frost CL. Reproductive parasitism: maternally inherited symbionts in a biparental world. Cold Spring Harb Perspect Biol. 2015;7(5):a017699.

Hussain, M., Akutse, K. S., Ravindran, K., Lin, Y., Bamisile, B. S., Qasim, M., Dash, C. K. and Wang, L. 2017, Effects of different temperature regimes on survival of *Diaphorina citri* and its endosymbiotic bacterial communities. Environ Microbiol, 19: 3439–3449.

Jeffries, C. L., and T. Walker 2016. Wolbachia Biocontrol Strategies for Arboviral Diseases and the Potential Influence of Resident *Wolbachia* Strains in Mosquitoes. Curr Trop Med Rep. 2016; 3: 20–25. doi:10.1007/s40475-016-0066-2

Jiggins FM 2017. The spread of Wolbachia through mosquito populations. PLoS Biol 15(6): e2002780.

Kikuchi Y, Tada A, Musolin DL, Hari N, Hosokawa T, Fujisaki K, Fukatsu T. 2016. Collapse of Insect Gut Symbiosis under Simulated Climate Change. mBio. 7:e01578-16. doi:10.1128/mBio.01578-16.

Kriesner P, Conner WR, Weeks AR, Turelli M, Hoffmann AA. 2016. Persistence of a Wolbachia infection frequency cline in *Drosophila melanogaster* and the possible role of reproductive dormancy. Evolution 70(5):979-97. doi:10.1111/evo.12923.

Martin, O. Y., Goodacre, S. L. (2009) Widespread infections by the bacterial endosymbiont Cardinium in arachnids. J. Arachnol. 37:106–108. doi:10.1636/SH08-05.1

Morag N, Klement E, Saroya Y, Lensky I, Gottlieb Y. Prevalence of the symbiont *Cardinium* in *Culicoides* (Diptera: Ceratopogonidae) vector species is associated with land surface temperature. FASEB J. 2012 26:4025–34.

Moreira LA, Iturbe-Ormaetxe I, Jeffery JA, Lu G, Pyke AT, Hedges LM, Rocha BC, Hall-Mendelin S, Day A, Riegler M, Hugo LE, Johnson KN, Kay BH, McGraw EA, van den Hurk AF, Ryan PA, O’Neill SL. (2009). A *Wolbachia* symbiont in *Aedes aegypti* limits infection with dengue, Chikungunya, and Plasmodium. Cell. 139:1268–78.

Morrow JL, Frommer M, Royer JE, Shearman DC, Riegler M. 2015. Wolbachia pseudogenes and low prevalence infections in tropical but not temperate Australian tephritid fruit flies: manifestations of lateral gene transfer and endosymbiont spillover? BMC Evolutionary Biology 15: 202.

Peel, M. C., Finlayson, B. L., and McMahon, T. A. 2007. Updated world map of the Köppen-Geiger climate classification. Hydrol. Earth Syst. Sci., 11, 1633-1644, doi:10.5194/hess-11-1633-2007.

Ross, P. A., Wiwatanaratanabutr, I., Axford, J. K., White, V. L., Endersby-Harshman, N. M., & Hoffmann, A. A. (2017). Wolbachia Infections in *Aedes aegypti* Differ Markedly in Their Response to Cyclical Heat Stress. PLoS Pathogens, 13(1), e1006006. doi:10.1371/journal.ppat.1006006

Weinert LA, Araujo-Jnr EV, Ahmed MZ, Welch JJ (2015). The incidence of bacterial endosymbionts in terrestrial arthropods. Proc Biol Sci. 282.

Wernegreen JJ. Mutualism meltdown in insects: Bacteria constrain thermal adaptation.Current opinion in microbiology. 15(3):255-262. doi:10.1016/j.mib.2012.02.001.

Werren JH, Windsor DM. *Wolbachia* infection frequencies in insects: evidence of a global equilibrium? Proceedings of the Royal Society B: Biological Sciences. 2000;267(1450):1277–1285.

Yee TW 2015. Vector Generalized Linear and Additive Models: With an implementation in R. New York, USA: Springer.

Yixin H. Ye, Alison M. Carrasco, Yi Dong, Carla M. Sgrò, and McGraw E. A. 2016. The Effect of Temperature on Wolbachia-Mediated Dengue Virus Blocking in Aedes aegypti. Am J Trop Med Hyg. 94(4): 812–819. doi:10.4269/ajtmh.15-0801.

Zchori-Fein, E. and K. Bourtzis (eds). 2011. Manipulative tenants - Bacteria associated with arthropods (CRC press).

