## Supplementary material for "*Wolbachia, Cardinium* and climate: an analysis of global data"

### Supplementary Information

#### Supplementary Figure Captions:

**Figure S1**:
Taxonomic breakdown of the screen database for arthropod groups classified as Mandibulata, i.e., Hexapoda (including insects), Crustacea and Myriapoda. Plots are shown for the number of screens (i.e., the number of populations sampled), and the total number of individuals screened. Ordinal or superordinal taxa are indicated if they comprise >2% of the database, on either measure. The full database is available as Supplementary Table S1.

**Figure S2**:
Taxonomic breakdown of the screen database for arthropod groups classified as Chelicerata, i.e., Acari (ticks and mites), Araneae (spiders), and relatives. All other details match Figure S1.

**Figure S3**:
The range of mean monthly temperatures for the period 1970-2010 associated with each of the Köppen climate classifications. These classifications are fit as a categorical predictor, along with mean temperature itself, in our analyses of symbiont prevalence (see Table 1; Supplementary Table S2). Bars show means, and confidence intervals show 95% quantiles. The upper row (panels (a) and (d)) shows all records in the WorldClim 2 database (Fick and Hijmans 2017). The middle row (panels (b) and (e)) shows the equivalent results for locations in our database of *Cardinium* screens, and the lower row (panels (c) and (f)) shows results for our database of *Wolbachia* screens. The left-hand column (panels (a)-(c)), shows results for the five higher-level climate classifications, namely A (tropical), B (arid), C (temperate), D (continental), and E (polar). The right-hand column (panels (d)-(f)): shows results for the 31 finer-grained climates, namely: Af (tropical rainforest), Am (tropical monsoon), Aw (tropical savanna, wet), As (tropical savanna, dry); BWh (hot desert), BWk (cold desert), BSh (hot steppe), BSk (cold steppe), Csa (hot-summer Mediterranean), Csb (warm-summer Mediterranean), Csc (cool-summer Mediterranean), Cwa (Monsoon-influenced humid subtropical), Cwb (subtropical highland or temperate oceanic climate with dry winters), Cwc (cold subtropical highland climate or subpolar oceanic climate with dry winters), Cfa (humid sub-tropical), Cfb (temperate oceanic), Cfc (subpolar oceanic); Dfa (hot-summer humid continental), Dfb (warm-summer humid continental), Dfc (subarctic), Dfd (extremely cold subarctic), Dwa (Monsoon-influenced hot-summer humid continental), Dwb (Monsoon-influenced warm-summer humid continental climate), Dwc (Monsoon-influenced subarctic climate), Dwd (Monsoon-influenced extremely cold subarctic climate); Dsa (hot, dry-summer continental), Dsb (warm, dry-summer continental), Dsc (sub-arctic, dry-summer continental), Dsd (extremely cold sub-arctic, dry-summer continental); ET (polar tundra), EF (eternal winter/ice cap). Comparing the rows shows that the some of the finer-grained climates were missing from our population sample (indicated by the gray labels), while others were represented by a biased sample with respect to temperature. For example, there were no population samples from EF (eternal winter/ice cap), while samples from ET (polar tundra) come from the warmer parts of this climatic zone.

**Figure S4:**
The effects of temperature on *Wolbachia* prevalence in the major insect orders (panels (b)-(g)), and in the remainder of the sampled Mandibulata (panel (h)). The slopes correspond to

results for three-parameter models given in Supplementary Table S2. All other details match Figure 2b-c, which is replicated for ease of comparison in panel (a).

**Figure S5**:
Best-fit slopes for the regression of mean *Wolbachia* prevalence onto temperature. Each point represents samples from a particular group of mandibulate hosts (one of the major insect orders, or the remainder of the data), and a single higher-level Köppen zone (polar: blue; continental: green, temperate: yellow, arid: orange; and tropical: red). Regressions that were individually significant are indicated as asterisks, and a full description is found in Supplementary Table S3. As expected, data show that the estimates of the slopes become more erratic as sample size decreases, but there is little evidence for a consistent trend towards negative or positive slopes. The apparent trend for *Wolbachia* infection to increase with temperature is driven by the well-sampled Diptera and Lepidoptera in temperate zones, indicated by the yellow asterisks towards the right-hand side of the plots.

**Figure S6**:
Best-fit mean prevalence estimates in Chelicerata hosts, for the symbionts *Cardinium* (left-hand bars) and *Wolbachia* (right-hand bars). In each case, estimates are shown for the five higher-level Köppen climates: from left-to-right, these are polar (blue), continental (green), temperate (yellow), arid (orange) and tropical (red).

**Figure S7**:
Estimates of mean temperature show strong correlations with other potential predictors of endosymbiont prevalence. Results are shown for (a) absolute latitude, (b) "seasonality" (defined as the standard deviation in the recorded temperatures over the period 1970-2000), (c) the maximum recorded temperature over this period, and (d) the minimum recorded temperature. All data come from the WorldClim database (Fick and Hijmans 2017).

**Figure S8:**
Best-fit mean prevalence values for *Cardinium* prevalence in Mandibulata hosts, in each of the Köppen climate zones. The upper row of panels ((a), (c) and (e)) shows estimates when only the climatic zones were included in the model, while the lower row (panels (b), (d) and (f)), show estimates after the effects of mean temperature have been removed. The strong difference between these rows suggests that temperature is a strong predictor of mean prevalence in these data. Results are compared for the five higher-level classifications (panels (a)-(d)), and for the 31 finer-grained classifications (panels (e)-(f)), not all of which were represented in our sample of screens (shown by the gray labels). Results are also compared when the model was fit only to mean prevalence (panels (a)-(b)), and when it was also fit the correlation parameter, allowing the shape of the distribution of prevalences to vary between climate zones (panels (c)-(d)). The definitions of all climatic zones, and the range of temperatures that they encompass, are shown in Supplementary Figure S3.

**Figure S9**
Best-fit mean prevalence values for *Wolbachia* prevalence in Mandibulata hosts, in each of the Köppen climate zones. The broad similarity between the upper and lower row of panels suggests that mean temperature is not a strong predictor of prevalence for these data. All other details match Supplementary Figure S8.

**Figure S10**:

Best-fit mean prevalence values for *Cardinium* prevalence in Chelicerata hosts, in each of the Köppen climate zones. The broad similarity between the upper and lower row of panels suggests that mean temperature is not a strong predictor of prevalence for these data. All other details match Supplementary Figure S8.

**Figure S11**:
Best-fit mean prevalence values for *Wolbachia* prevalence in Chelicerata hosts, in each of the Köppen climate zones. The broad similarity between the upper and lower row of panels suggests that mean temperature is not a strong predictor of prevalence for these data. All other details match Supplementary Figure S8.

**Supplementary Tables:**

**Supplementary Table S1**: The database of screens of wild arthropod populations for *Cardinium* or *Wolbachia*.

**Supplementary Table S2**: results of Beta-binomial model fitting to symbiont screen data.

**Supplementary Table S3**: tests for an effect of mean temperature of *Wolbachia* prevalence in Mandibulata hosts, applied separately to populations from each of the five higher-level Köppen climates.

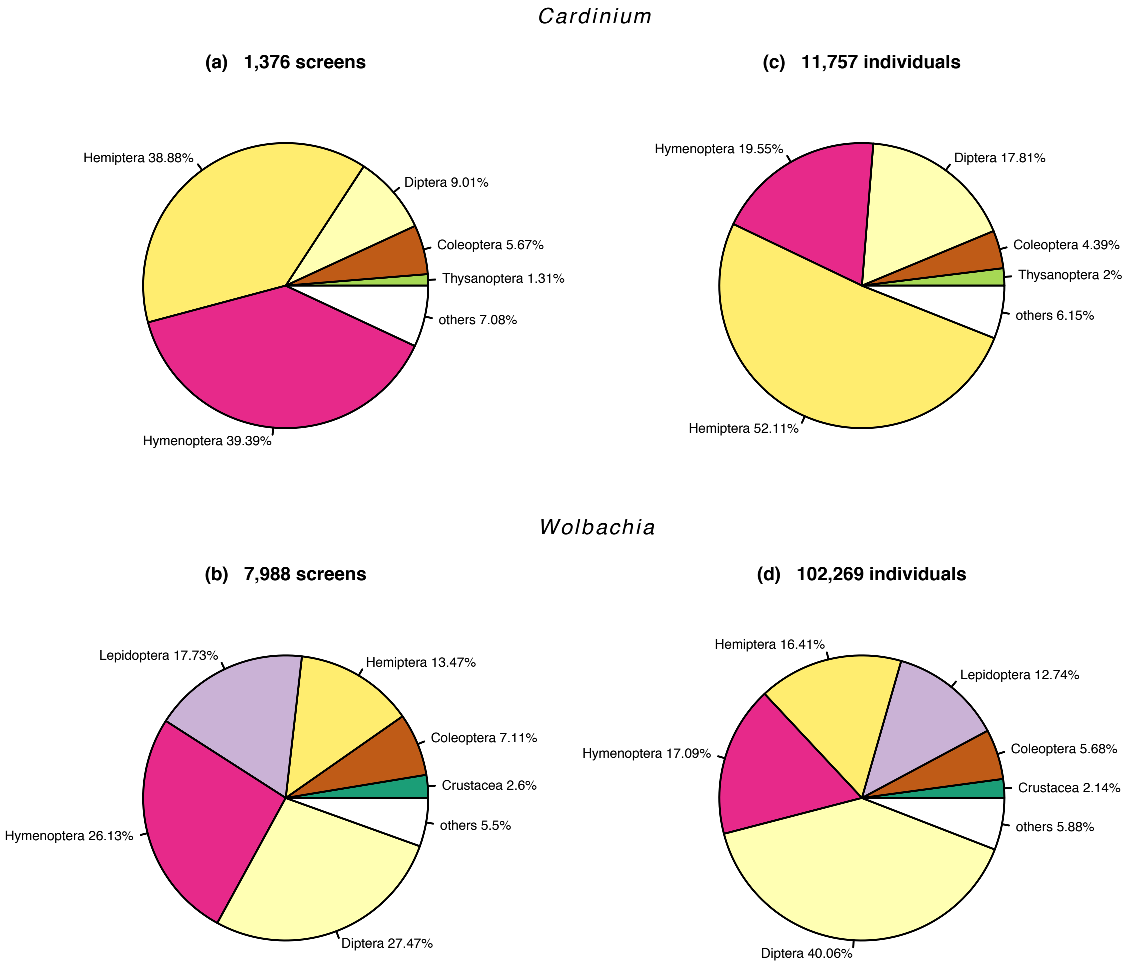
Cardinium
(a) 1,376 screens
Hemiptera 38.88%
Diptera 9.01%
Coleoptera 5.67%
Thysanoptera 1.31%
others 7.08%
Hymenoptera 39.39%
(c) 11,757 individuals
Hymenoptera 19.55%
Diptera 17.81%
Coleoptera 4.39%
Thysanoptera 2%
others 6.15%
Hemiptera 52.11%
Wolbachia
(b) 7,988 screens
Lepidoptera 17.73%
Hemiptera 13.47%
Coleoptera 7.11%
Crustacea 2.6%
others 5.5%
Hymenoptera 26.13%
Diptera 27.47%
(d) 102,269 individuals
Hemiptera 16.41%
Lepidoptera 12.74%
Hymenoptera 17.09%
Coleoptera 5.68%
Crustacea 2.14%
others 5.88%
Diptera 40.06%

Figure S1.

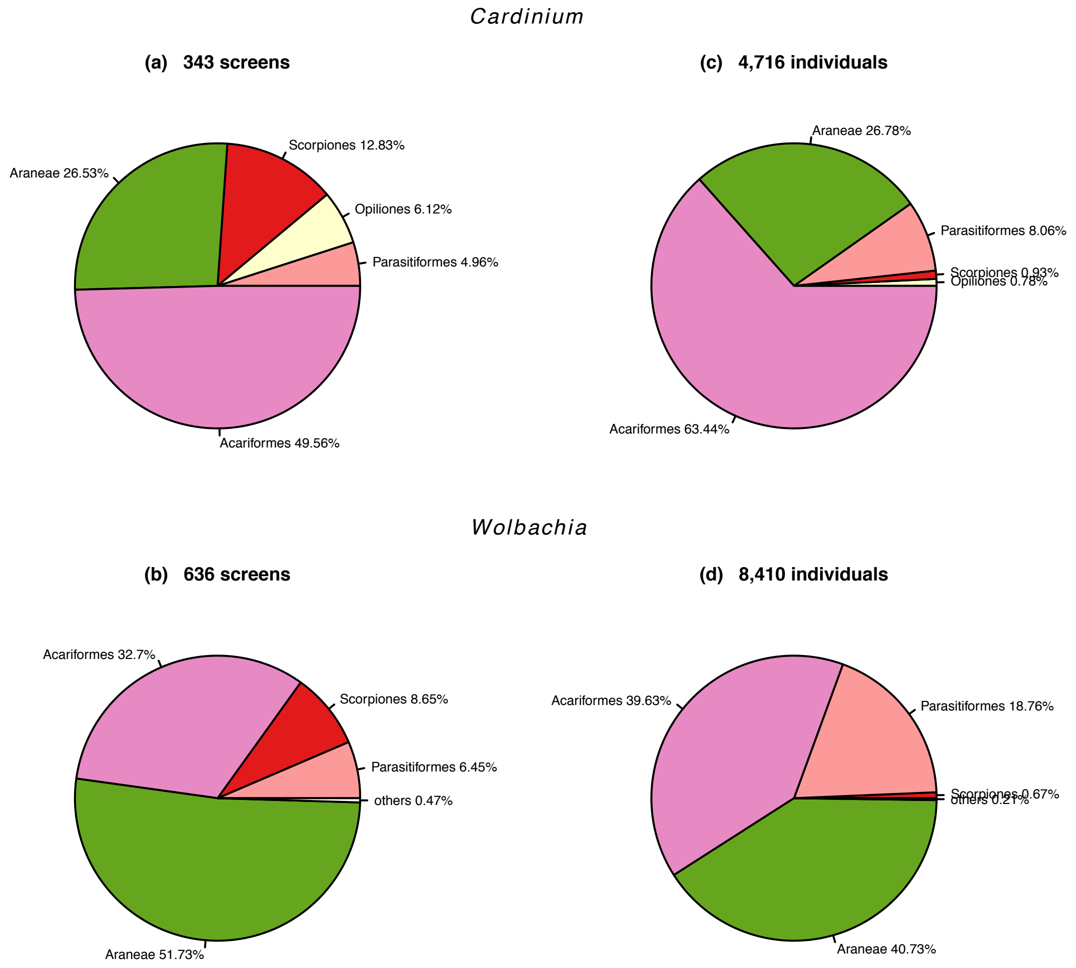
Cardinium
(a) 343 screens
Scorpiones 12.83%
Araneae 26.53%
Opiliones 6.12%
Parasitiformes 4.96%
Acariformes 49.56%
(c) 4,716 individuals
Araneae 26.78%
Parasitiformes 8.06%
Scorpiones 0.93%
Opiliones 0.78%
Acariformes 63.44%
Wolbachia
(b) 636 screens
Acariformes 32.7%
Scorpiones 8.65%
Parasitiformes 6.45%
others 0.47%
Araneae 51.73%
(d) 8,410 individuals
Acariformes 39.63%
Parasitiformes 18.76%
Scorpiones 0.67%
others 0.21%
Araneae 40.73%

Figure S2.

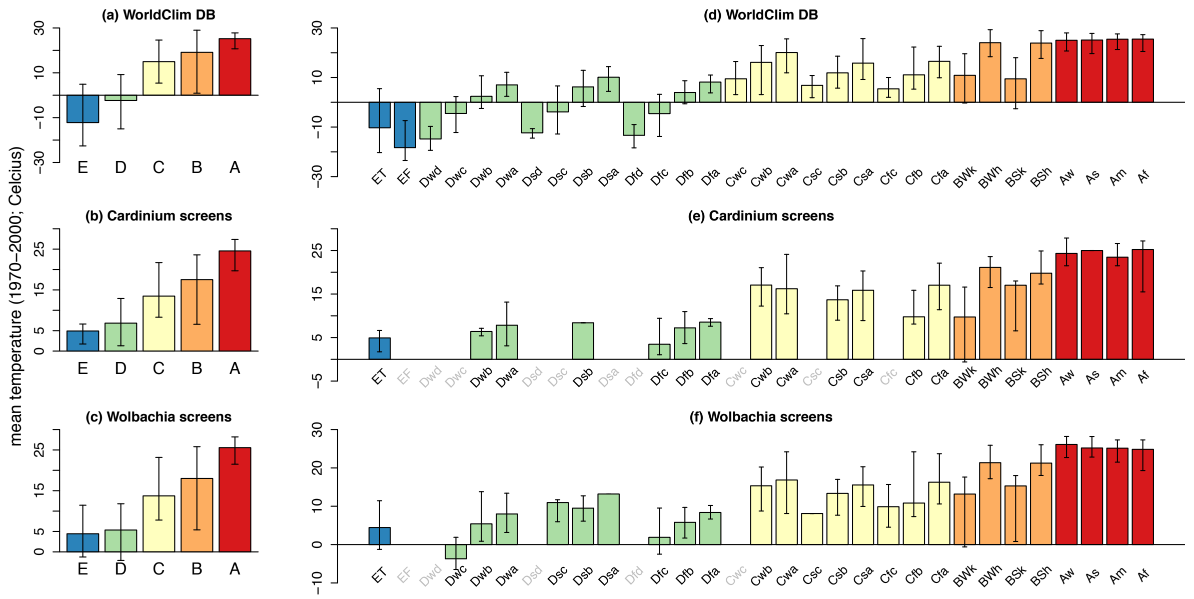


Figure S3.

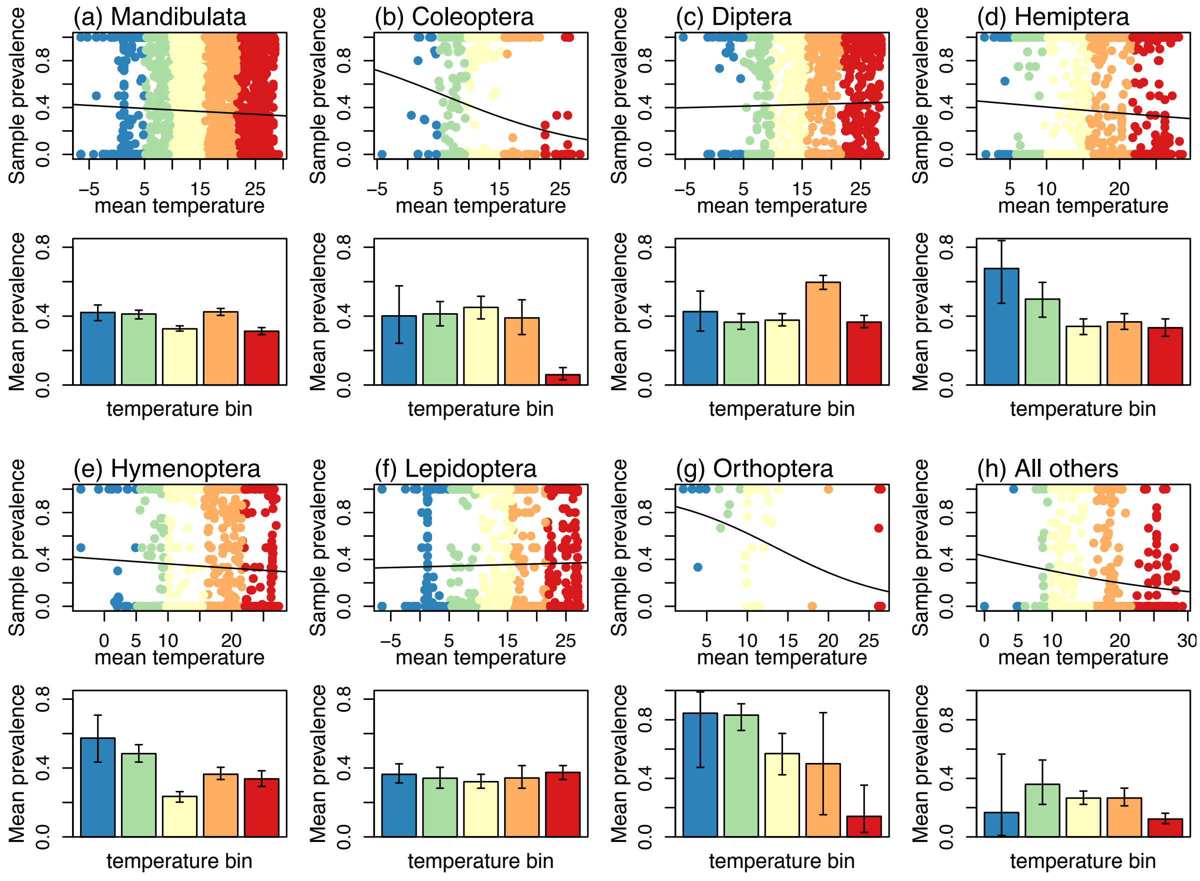


Figure S4.

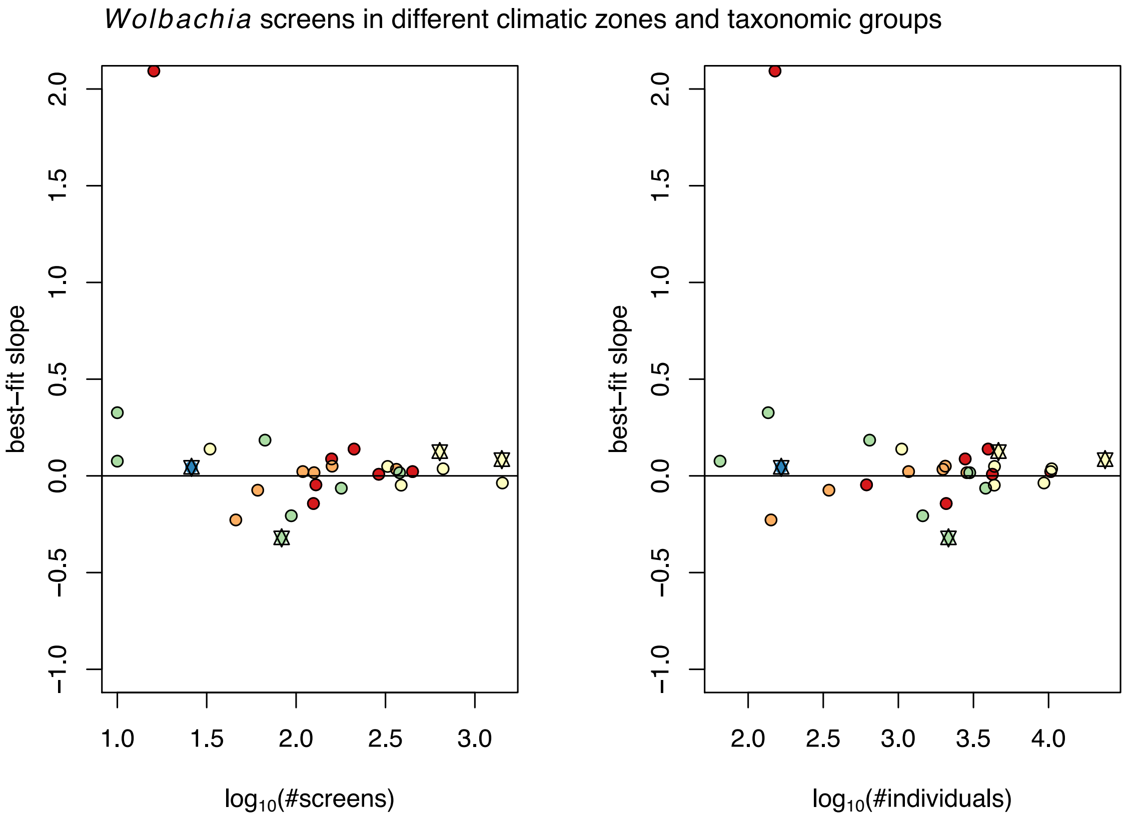


Figure S5.

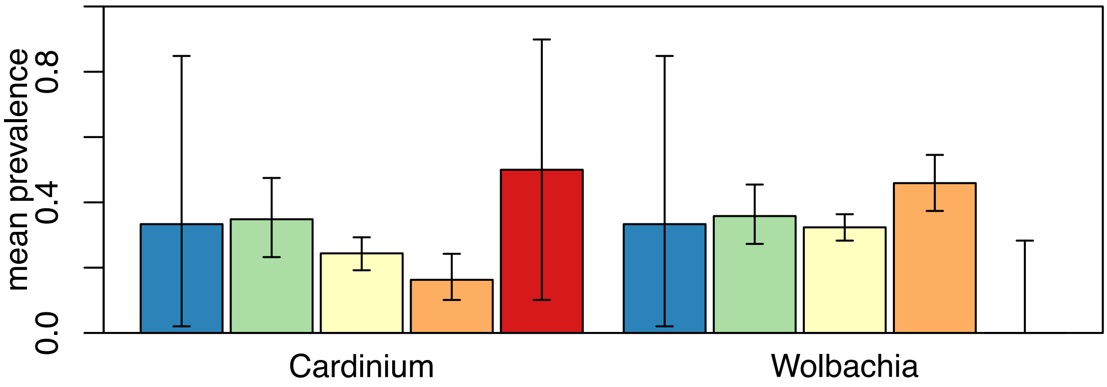


Figure S6.

**Correlations between WorldClim bioclimatic temperature variables**

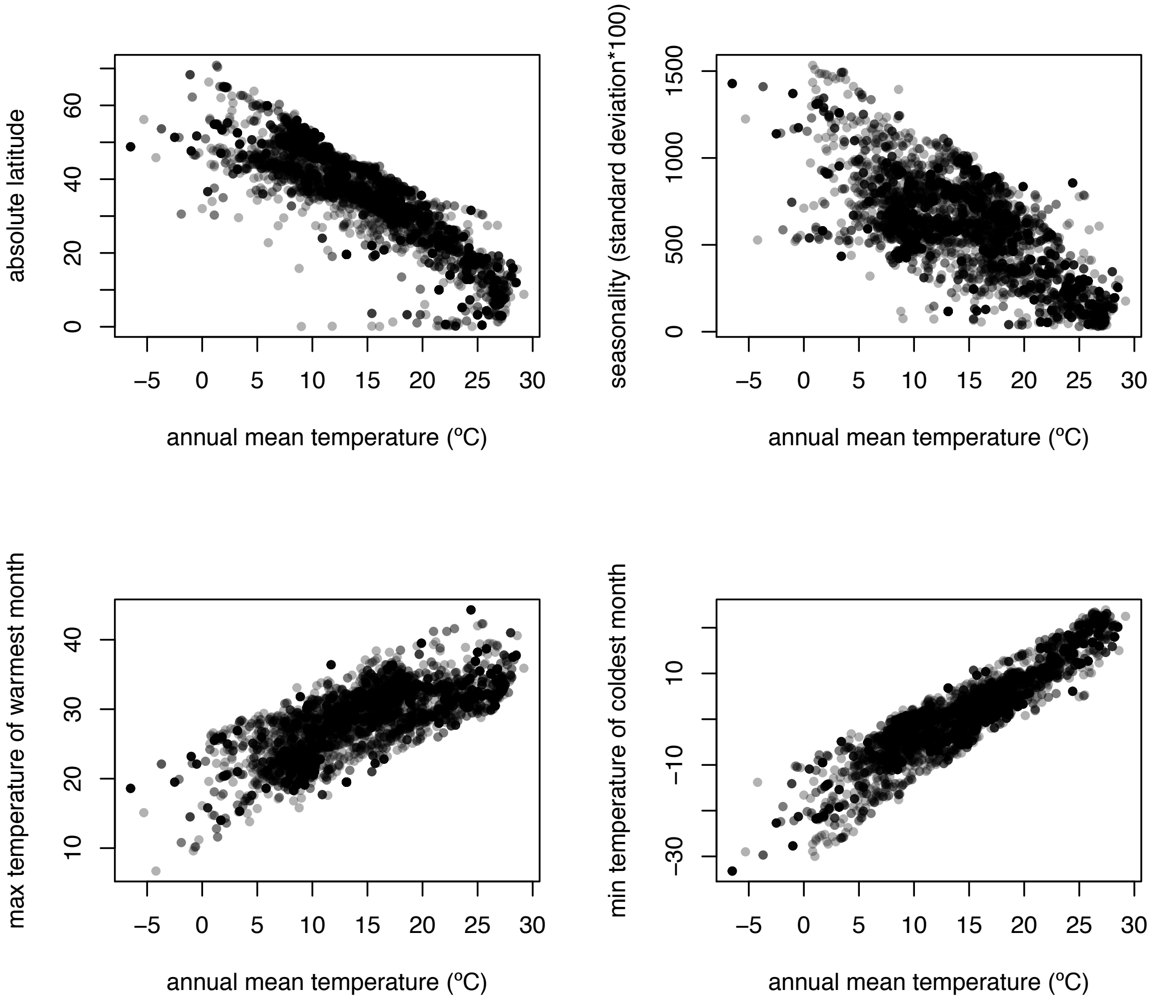


Figure S7.

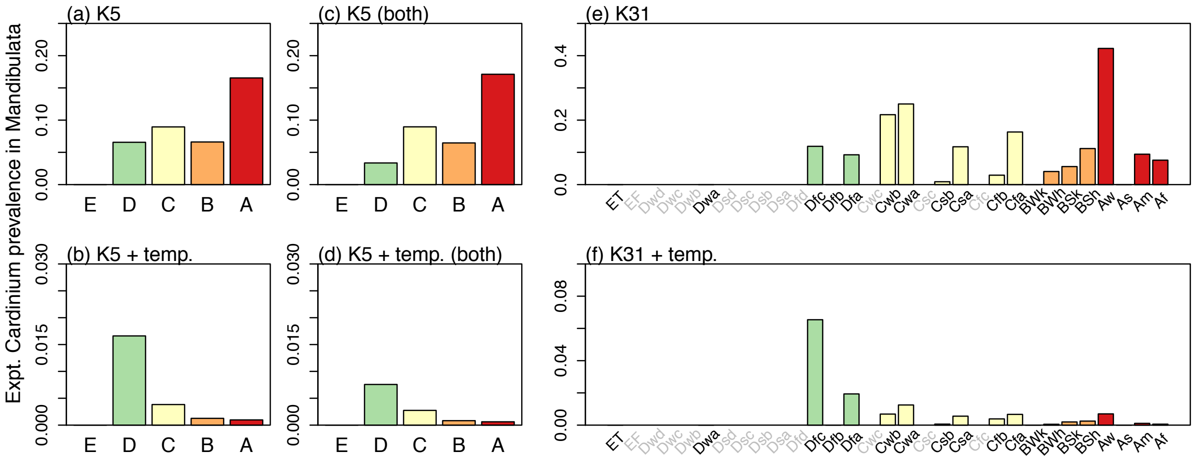


Figure S8.

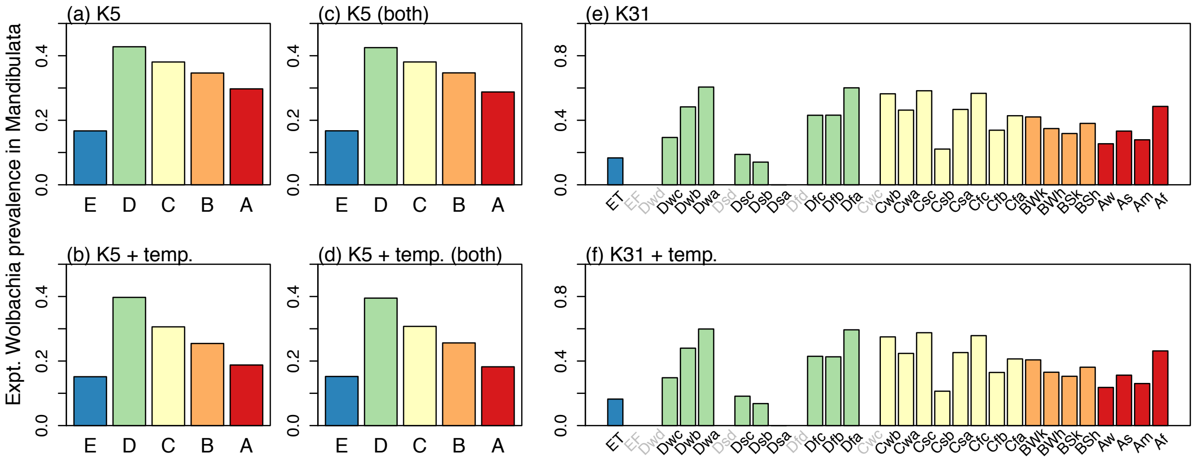


Figure S9.

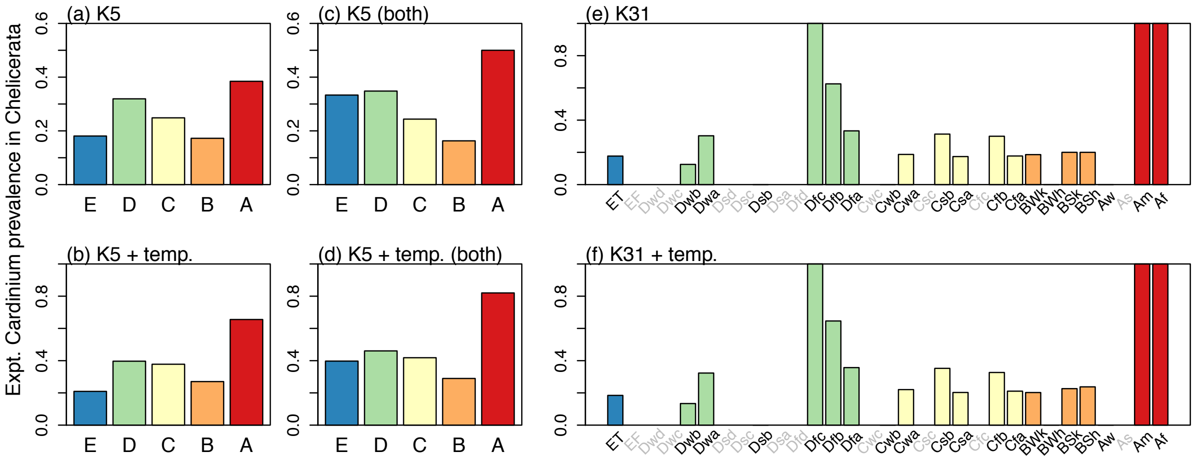


Figure S10.

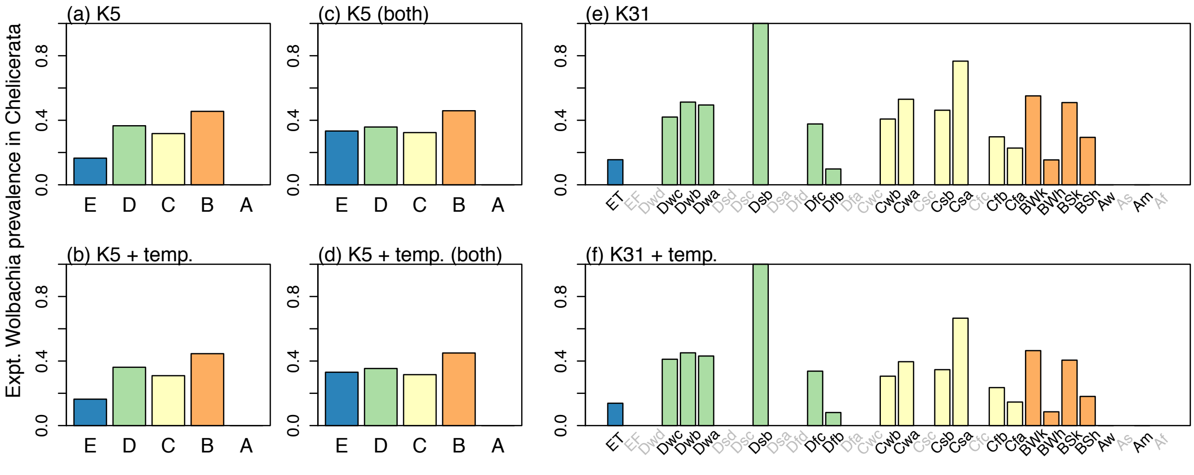


Figure S11.
